# Web-based eye-tracking for remote cognitive assessments: The anti-saccade task as a case study

**DOI:** 10.1101/2023.07.11.548447

**Authors:** Gustavo E Juantorena, Francisco Figari, Agustín Petroni, Juan E Kamienkowski

**Author notes:** Equal contributions (Gustavo E Juantorena, Francisco Figari).

## Abstract

Over the last years, several developments of remote webcam-based eye tracking (ET) prototypes have emerged, testing their feasibility and potential for web-based experiments. This growing interest is mainly explained by the possibility to perform tasks remotely, which allows the study of larger and hard-to-reach populations and potential applications in telemedicine. Nevertheless, a decrease in the quality of the camera and a noisier environment bring new implementation challenges. In this study, we present a new prototype of remote webcam-based ET. First, we introduced improvements to the state-of-the-art remote ET prototypes for cognitive and clinical tasks, e.g. without the necessity of constant mouse interactions. Second, we assessed its spatiotemporal resolution and its reliability within an experiment. Third, we ran a classical experiment, the anti-saccade task, to assess its functionality and limitations. This cognitive test compares horizontal eye movements toward (pro-saccades) or away from (anti-saccades) a target, as a measure of inhibitory control. Our results replicated previous findings obtained with high-quality laboratory ETs. Briefly, higher error rates in anti-saccades compared to pro-saccades were observed, and incorrect responses presented faster reaction times. Our web-ET prototype showed a stable calibration over time and performed well in a classic cognitive experiment. Finally, we discussed the potential of this prototype for clinical applications and its limitations for experimental use.

## Introduction

Eye movements are an insightful window to cognitive and emotional functioning. By measuring gaze position on a screen with an eye tracker, researchers can infer and study cognitive processes in different settings and tasks (Hayhoe and Ballard 2005; Just and Carpenter 1980; Linari et al. 2022; Liversedge and Findlay 2000; Rayner 1998; Schütz, Braun, and Gegenfurtner 2011). Some important advantages of eye tracking (ET) compared to other tools to study human cognition are i) its non-invasiveness and absence of body sensors ii) relative low cost, in the range of 10-50 thousand USD for research level ET iii) relative portability, compared to functional magnetic resonance (fMRI), magnetoencephalography (MEG) or even electroencephalography (EEG) and iv) good temporal resolution, in the scale of milliseconds for research equipment.

Over the last years, different developments and types of ET devices have emerged. For practical purposes, we will group ET devices in three main categories, 1) High-resolution standard laboratory equipment 2) High portability, lower resolution ET and 3) Webcam based, low-resolution ET. The first group, with high temporal and spatial resolutions like EyeLink, usually has the downside of a higher cost and poor usability outside the lab. The second group comprises more flexible and portable equipment, but with lower temporal resolutions and similar costs (e.g. Tobii). Within this group, some cheaper developments appeared in the market, like Gazepoint and EyeTribe, which are more oriented to marketing applications. The third group, which includes the prototype we present in this study, is a more recently explored alternative that consists on performing ET in domestic computers using standard webcams, removing the need of specialized hardware (Huang et al. 2016; Hutt et al. 2023; Papoutsaki et al. 2016; Semmelmann and Weigelt 2018; Slim and Hartsuiker 2022; vanWell and Tanaka 2022; Xu et al. 2015; Yang and Krajbich 2020). With this technology, there is no need to attend a laboratory, neither the supervision of an experimenter (in-person or online), and can be performed simultaneously by a large number of participants. These developments imply that any standard laptop or desktop computer with a webcam can perform ET, bringing ET to any potential participant with a standard home computer, and opening potential web applications and its use in telemedicine. Of course, this type of ET system has important limitations, like lower frame rate, lower spatial resolution, and less accuracy. Nevertheless, some simple cognitive ET tasks do not require high resolution, being potential good candidates for these setups.

In the case of web-based ET cognitive experiments, its asynchronous nature implies that no technician or experimenter needs to be present while the subject performs the task. This advantage, in turn, imposes additional conditions. First, clear instructions, transmitted solely through text, images or videos, must be provided. Second, as in other remote experiments, the concentration of the participant will be lower, so the experiments must have a lower number of trials, more attentional cues, and catch trials. And third, there must be some remote evaluation of the environment, e.g., light conditions, quality of the hosting hardware (Huang et al. 2016), head movements, and distance of the participant to the screen (Hansen and Ji 2010; Li et al. 2020). The two first conditions can be addressed with a better experimental design of the cognitive task, while the third condition impose technical constraints that need to be addressed by design improvements of the web-based ET. For instance, Xu and collaborators measured light conditions and, if needed, requested the subject to adjust the ambient light (Xu et al. 2015). They also verified that a minimum frame rate was satisfied before starting the experiment. Papoutsaki and colleagues requested the subjects to center their heads with respect to the webcam (Papoutsaki et al. 2016). Additionally, Li and colleagues proposed a method to estimate the screen size (in cm) and the distance to the screen using objects of standard size and the participant’s blind spot location (Li et al. 2020). We adapted this tool for jsPsych (https://www.jspsych.org/) (de Leeuw 2015) in collaboration with Peter J. Kohler and Josh de Leeuw.

Another important limitation when using web-based ET is the gaze estimation algorithms that can be utilized. Given the constraint of having access to only one camera, a small fraction of the gaze estimation algorithms found in the bibliography can be used, namely appearance-based methods (Hansen and Ji 2010; Xu et al. 2015). But none of these algorithms guarantee invariance toward head movements (i.e., the capacity of the system to keep working despite the subject moving his head) (Hansen and Ji 2010). Since head movements cannot be restricted, frequent decalibrations are expected. Thus, frequent recalibrations of the system are needed, which map the images of the eyes to coordinates on the screen. On the one hand, Xu et al. (Xu et al. 2015) presented a series of stimuli on the calibration screen, collecting several frames per stimulus. From this sequence of frames, they discarded those that differ notoriously from the average, as they assumed they corresponded to blinks, and kept the last frames for each stimulus presentation. On the other hand, both Huang et al. (2016) and Papoutsaki et al. (2016), collected one single frame per user interaction (e.g., clicks, typing). The former analyzed at which instant the gaze better coincides with the coordinate of the interaction for different types of interactions and selected the frame based on this criteria. The latter simply uses the frame immediately prior to the interaction as they assume that the gaze arrives before the hand (mouse pointer) to a visual target.

One of the most promising applications of web-based ET is in the field of cognitive sciences and potentially for neuropsychiatric evaluations or telemedicine. To make progress in this direction, research efforts should be first directed toward replicating classic laboratory ET experiments. Semmelmann and colleagues (2018) compared three typical eye-movement patterns performed in a cognitive task recorded in a web-based eye tracker with a laboratory eye tracker (Semmelmann and Weigelt 2018). Although they didn’t replicate a particular cognitive task, they obtained similar results to those obtained with a high-resolution ET. In 2022, Slim and Hartsuiker replicated a cognitive task in a web-ET called visual world study that was previously conducted by Dijkgraaf and colleagues (Dijkgraaf, Hartsuiker, and Duyck 2017; Slim and Hartsuiker 2022). Bánki and colleagues (2022) attempted to replicate lab results with web-based ET in infants, using an audio-visual synchrony perception task (Bánki et al. 2022). They could not replicate the lab results to their full extent, most probably because of the challenges associated with infants, since they perform more body and head movements than adults. Nevertheless, the system could not be confronted given that it is closed and not published (LabVanced). Yang and Krajbich (2021) used an open ET library called WebGazer (Papoutsaki et al. 2016) integrated with jsPsych and performed a binary choice task, replicating previous lab results (Yang and Krajbich 2020). Madsen and colleagues (2021) studied the engagement of students in online education in a remote setting, with a web-based ET (Madsen et al. 2021). They found that synchronization of eye movements among students is a good predictor of individual learning performance.

In this study, we describe a prototype of an ET device that presented a good performance in a classic cognitive experiment. The prototype was tested on different computers, cameras and browsers using a simple calibration-like experiment. Then, we tested the prototype running the anti-saccade task (Currie et al. 1991; Hallett 1978; Hutton and Ettinger 2006; Munoz and Everling 2004). In this task, the participant is first instructed to fixate their gaze on a central cue, which indicates whether they have to look toward a lateralized cue (pro-saccades) or in the opposite direction, away from the lateral cue (anti-saccades). After a short interval fixating the central cue, the lateralized cue appears, and the saccade latency and the error rate is computed for both pro- and anti-saccades. The anti-saccade task measures the capacity of an individual to inhibit the automatic gaze movement toward the appearing lateral visual cue, to instead perform a voluntary saccade in the opposite direction (Munoz and Everling 2004). Depending on the design of the task, working memory and visuo-spatial memory can also be studied (Olincy et al. 1997; Unsworth et al. 2011; Unsworth, Miller, and Robison 2021).

The anti-saccade task is very simple and has clear expected results that are fairly well established. It has been studied in individuals with low working memory capacity (Unsworth et al. 2011) and in relation to age (Munoz and Everling 2004; Olincy et al. 1997; Płomecka et al. 2020). Additionally, it is sensitive to a broad range of neurological populations (Everling and Fischer 1998; Hutton and Ettinger 2006), and it was first studied in frontal lobe lesions (Guitton, Buchtel, and Douglas 1985). Thus, it is a good candidate for a validation case study and a first step in the use of remote eye trackers in patient populations.

Here, we propose several improvements to state-of-the-art web-based ET systems. Starting from existing implementations, we built a new prototype of a web-based eye tracker. This system is intended to not be tied to the present case study but to have a flexible implementation that could be used for other short tasks, including those that do not use mouse responses. We were able to replicate the results of the anti-saccade task, paving the way for a large number of applications, including cognitive testing in patient populations. Finally, we reflect on the intrinsic limitations of the web-based ET for future developments.

## Methods

### Eye-tracker prototype development

#### FACEMESH

An eye-tracking (ET) device provides an estimation of the gaze position on the screen. In order to do this, it is important to first solve the subproblem of the eyes’ localization in space (Hansen and Ji 2010). That is, where the eyes of the user are located given a series of frames captured by the camera. After a revision of the state-of-the-art alternatives, WebGazer was used as a starting point for our prototype, which already proposed a solution to both problems: gaze position and eyes location.

Eyes localization was solved by using a facemesh model and the frames of the webcam as its input. This model generates, in real time, a 3D reconstruction of the user’s face as output. The reconstruction is represented in a mesh (i.e., net of interconnected spatial nodes) in which each node maps to a specific position of the face. With this mesh, queries can then be done on each frame about the appearance of the user’s face (for example, to extract a couple of rectangles that enclose each eye).

The original facemesh model used by WebGazer crashed after around 10 minutes ^1^. To solve this, we replaced the original facemesh package which at the time was already deprecated (@tensorflow-models/facemesh/v/0.0.3 to @tensorflow-models/face-landmarks-detection /v/0.0.3). TensorFlowJS’ version was also put up to date (@tensorflow/tfjs/v/2.0.1 to @tensorflow/tfjs/v/3.13.0). While doing that, modifications were also done to the section of code which handled the retrieval of the eye coordinates ^2^. These specific changes were later merged to the original WebGazer repo though not through a pull request ^3^. When working with this code it is recommended to keep the different modules up to date.

#### CALIBRATION

Gaze estimation is solved by applying a Ridge Regression model that uses the appearance of the localized eyes as the independent variable and the expected position as the dependent variable. Thus, for the input, each rectangle is first resized to a fixed size of 10 x 6 pixels, and converted to grayscale to which histogram equalization is applied. Both rectangles are then transformed into a single input vector of 120 dimensions. The dependent variable is a pair of coordinates corresponding to the expected gaze position of the user on the screen. Then, after fitting the model, it is possible to transform the images constantly captured by the camera into a pair of coordinates on the screen corresponding to the estimated gaze position of the user. This implementation of gaze estimation is the one originally chosen in the WebGazer package and remained unchanged.

To instantiate the gaze estimation model, a series of pairs <eyes images, screen coordinates> is required. In the WebGazer project, this input data is collected continuously after clicks and mouse movements (Papoutsaki et al. 2016). This decision is based on the assumption that gaze position and pointer position are synchronized when the user clicks or moves the mouse over the screen. This hypothesis and its limitations had previously been studied by Huang and colleagues (Huang et al. 2016).

However, our task did not include mouse responses from the user and thus this type of calibration was inadequate. Instead, we implemented explicit calibration phases in which calibration points are shown to the user in specific locations. For each of them, the user had to fixate the gaze on the dot while pressing the spacebar at the same time, similar to traditional eye trackers. At this point, another optimization was done over WebGazer. Since continuous calibration data was originally allowed, the coefficients of the regression model were computed on each frame demanding constant computational efforts. With the explicit calibration phases, these coefficients only need to be computed once at the end of each of these phases and thus the implementation was adjusted accordingly ^4^.

#### HEAD MOVEMENTS

The gaze estimation strongly depends on the head position. Given the impossibility of restricting the movement of the user and the lack of head movement invariance, we also implemented a notification of recalibration suggestion which is based on detecting movements from the user. During calibration, the positions of the eyes of the user are stored. Then, on each frame, a check is done to verify if the eyes are in a similar position to the one established during the calibration phase. If a movement is detected, an event is emitted which has to be interpreted as a suggestion for the tool to be recalibrated.

For this to be possible, an extra improvement was done to the WebGazer package. Originally the computed coordinates of the localized eyes were private to the package and thus only visible from inside its running script. To allow its reuse, these variables were instead exposed through JS custom events, as they would then be consumed by our head movement detection module ^5^.

#### INTERFACES & PLAYGROUNDS

Compatibility with jsPsych (de Leeuw 2015) had to be provided which is why some code is provided to facilitate the writing of jsPsych timelines. At the same time we wanted to keep the eye tracker accessible directly from the browser, as to allow users of the system outside of jsPsych’s context. These two use cases imply each a different interface with which the system is accessed.

For each interface a playground was implemented. These are a couple of minimal HTML and JS files that allow interaction with the features provided by each interface. By providing a minimal usage scenario, they facilitate the development of each module. These are accessible in the repository’s homepage (Please refer to *Code availability* section).

### Virtual chin-rest

The virtual chinrest (Li et al. 2020) is a method that measures the screen resolution and viewing distance of the participant to the screen in a web browser. This allows automatic adjustment of stimulus settings based on the individual’s viewing distance. In summary, the virtual chinrest consists of two steps: Firstly, the screen resolution was estimated by asking the participant to put a card (credit, debit, id, etc) on the screen and adjust a frame by dragging one of its corners to match it to the size of the screen (Fig. 1.A.I). As these cards had a standard size (85×55 mm) it is possible to convert pixels (of the frame) to millimeters (of the card). This makes it possible to calculate the logical pixel density (LPD) (in pixels per mm, Eq. 1). Secondly, the distance of the participant to the screen was estimated by performing a simple experiment to estimate their blind-spot. The participant was asked to cover their right eye and focus on a small black square on the right side of the screen. After pressing the bar, a small red dot starts moving horizontally from the black square toward the left side of the screen (Fig. 1.A.II). The participant had to press the bar again when the red dot disappeared (into the blind-spot) as fast as possible. This procedure is repeated three times and their results were averaged. Then, based on the relatively conserved position of the blind spot location (α = 13.5°) and the distance from the blind-spot to the center of vision (Fig. 1.B), the viewing distance was calculated as eq. 2.

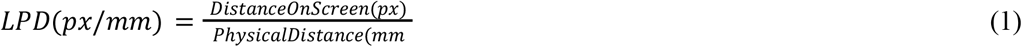

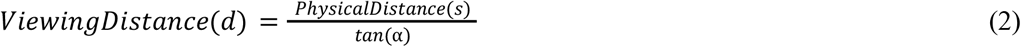

**Figure 1.**
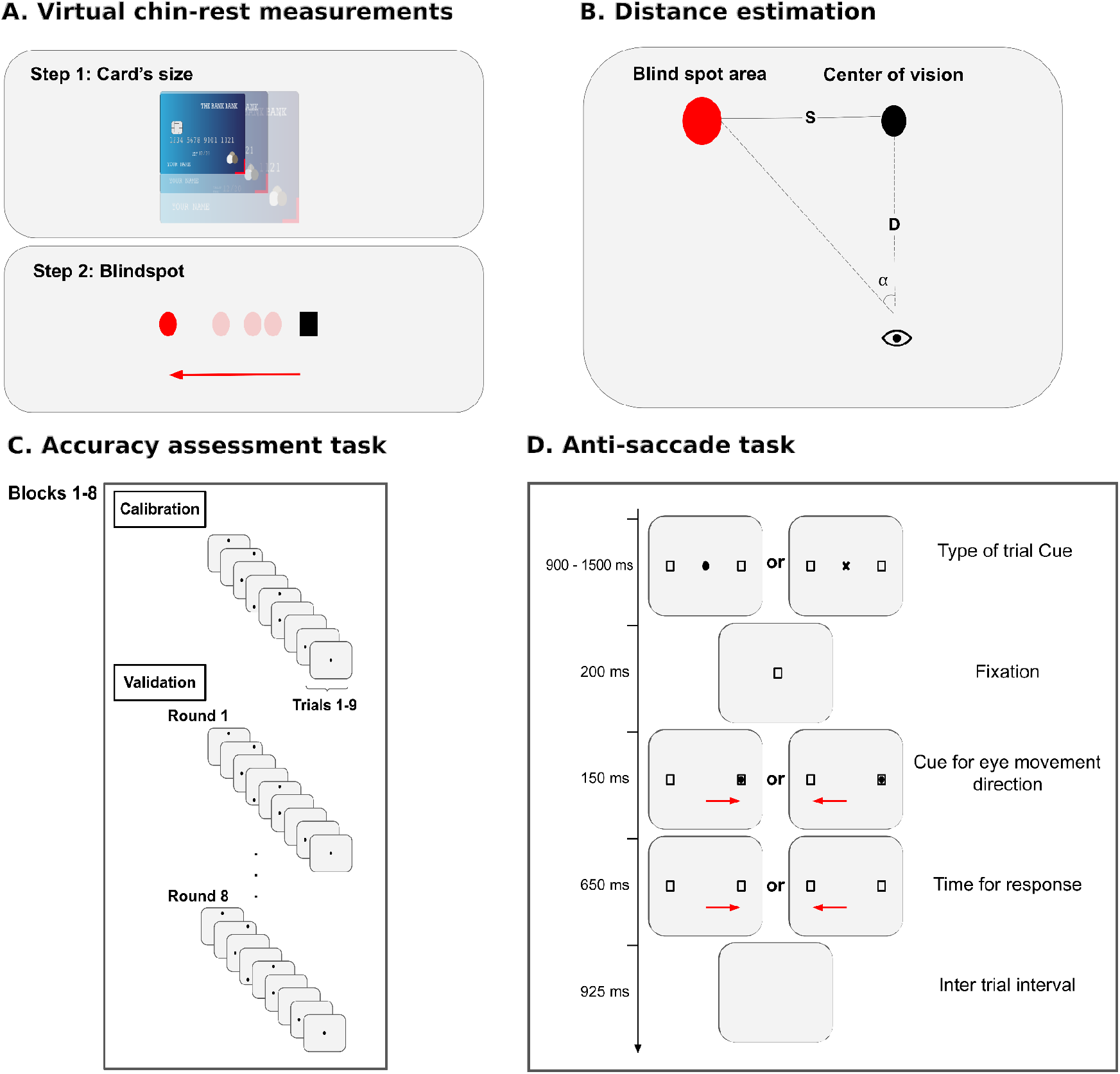
Protocols. **A-B) Virtual chin-rest.** Step 1: Measure by placing the card on screen and dragging the corner until sizes match. Step 2: Distance estimation by measuring the angle between the center of vision and the blind spot. The participant covers one eye and follows the red ball moving from left to right, touching the space bar when he loses sight of it (see equations (1) and (2)). **C) Accuracy assessment of the system**. The prediction error of the prototype is quantified based on point presentation. The structure of the task consists of 8 blocks. Each block has a 9-point calibration and then 8 validation rounds of 9 points each (trials). Finally, the error is evaluated in these 576 trials. **D) Anti-saccade task**. The task begins with the fixation of a central stimulus that is informative about the type of trial (cross for anti-saccade, empty square for pro-saccade). Then a cue appears briefly and when it disappears the participant must look to the corresponding side according to the type of trial.

This distance must be approximately corroborated by the participant, if not, the procedure is repeated.

We contributed to its development as a plugin for jsPsych^6^, and it has been included in the current version^7^. This allows us to present stimuli at a fixed size for different monitors and also returns the conversion factor from pixels to visual degrees (i.e., the visual angle of a single pixel), which is essential for converting gaze coordinates from pixels to visual angle degrees.

### Case study 1: Accuracy assessment

The initial assessment of the prototype’s performance with regards to reliability and temporal and spatial resolution was done by alternating calibrations (as described before) and validation stimuli. The validation stimuli were black dots that appeared in the same 9 positions of the calibration but in a random sequence. These stimuli allowed us to estimate the accuracy of the system (as participants are instructed to fixate in those positions) and the reliability (as the procedure was repeated several times). Also, during this experiment, we could explore the distribution of the sampling rate over time.

### Experimental protocol

In order to assess the quality of the gaze estimation of our system and its degradation over time, we proposed a protocol composed of the following stages: data loading of the hardware and software used, estimation of the distance to the screen using the virtual chinrest (Li et al. 2020), and evaluation of the prototype. The latter, in turn, consisted of eight trials in which the first calibration was presented where the subject must observe nine circles one at a time, wait for them to change from blue to black and, when this happens, press the spacebar. Only by pressing the bar, the circle disappeared and the next one appeared. Then, the validation period began, where eight trials were presented, consisting of a fixation cross in the center of the screen for 1800 milliseconds and then nine black circles identical to the previous ones. At this stage it was not necessary to perform any mouse or keyboard interaction, just fixate on them, and each one disappeared after 800 milliseconds. The circles in both stages were 20 pixels in radius and their presentation was always one at a time in a random fashion, within a predetermined grid at the center of the screen. Eight blocks were presented, with eight validation rounds per block of nine points each, for a total of 576 trials.

### Participants

Two of the authors completed the accuracy assessment (JEK: 43 y.o.; GEJ: 33 y.o.). Both have normal vision. The accuracy assessment was reviewed and approved by the IRB of Instituto de Investigaciones Médicas Alfredo Lanari (IDIM). All participants provided written informed consent in agreement with the Helsinki Declaration.

### Materials

In order to compare between cameras and remote versus local implementation, we used the same computer configuration (Ubuntu 20.04.5 LTS / 16 GB RAM / Intel® Core™ i5-4460 CPU @ 3.20GHz × 4) and screen (LG 22MP48HQ-P,1920X1080, 60 Hz). This is a standard configuration for nowadays computers. We also selected some standard cameras (see Table 1). The experiment was presented in the Chrome browser because it is the most popular in the world^8^.

**Table 1.**
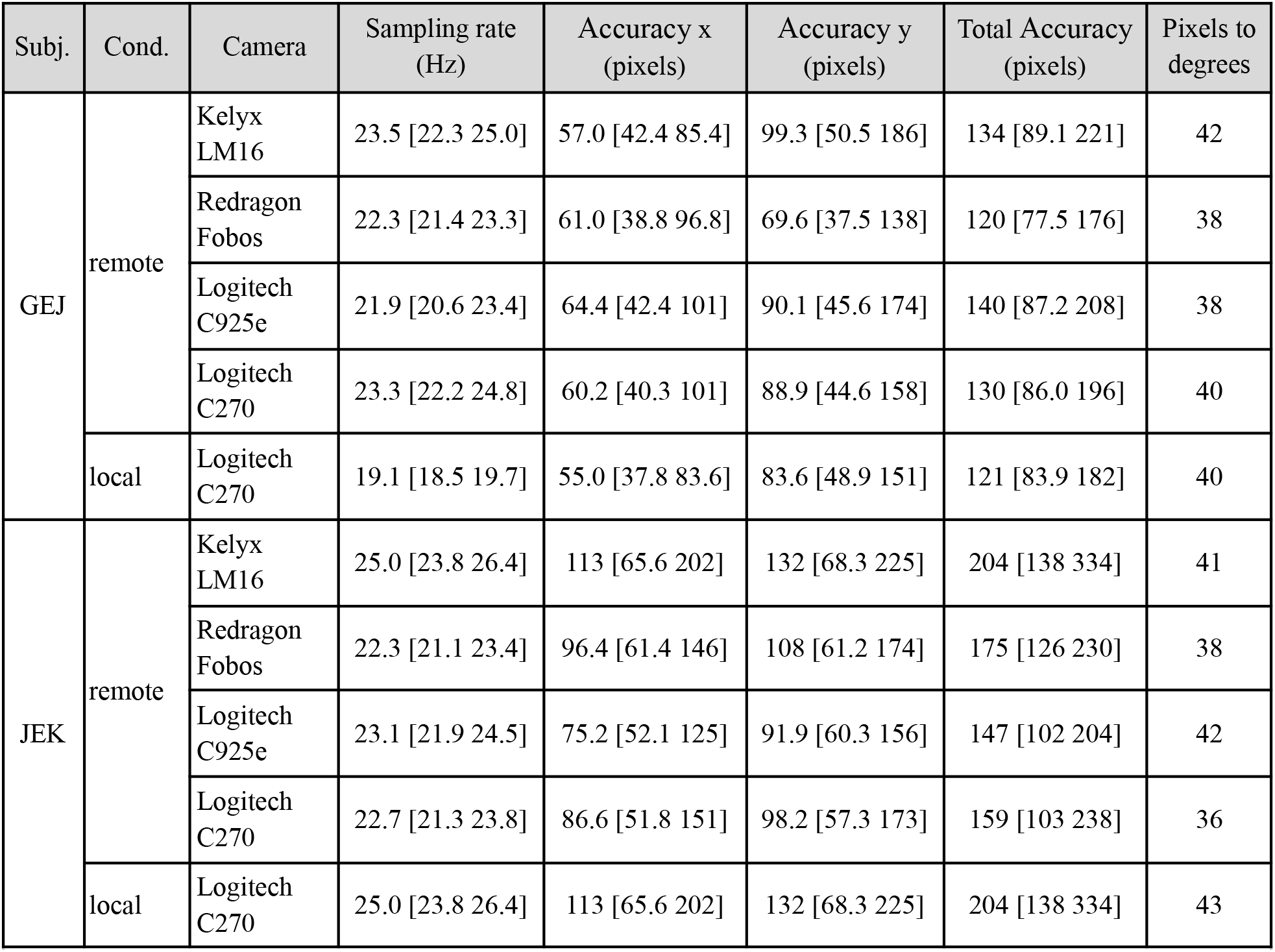
Accuracy assessment. Summary measures for every camera and condition for both experimental subjects. Values are expressed as median [IQR]. All cameras have a frame rate of 30 FPS. The resolution of the cameras are 720p (Logitech C270 and Redragon Fobos) or 1080p (Kelyx LM16 or Logitech C925e).

## Case study 2: The anti-saccade task

The anti-saccade task is a cognitive task with high clinical relevance. It has been used for many decades (Currie et al. 1991; Hallett 1978; Hutton and Ettinger 2006) and assesses the ability of an individual to inhibit prepotent responses, particularly those involving eye movements. This task requires a participant to look away from a visual target, either to the left or right, rather than toward it. The task typically involves the presentation of a target stimulus in one location, followed by a cue indicating the location to which the participant should make a saccade in the opposite direction of the target. The task requires the participant to suppress the natural tendency to look toward the target and instead make a saccade in the opposite direction, which is a cognitively demanding process.

### Experimental protocol

At the beginning of the protocol, each participant had to fill out a short form, allow access to their webcam, and complete the two virtual chinrest tasks (credit card and blind spot, see Virtual Chinrest). They were then shown the video captured by their webcam and asked to center their head in a box on the screen. In this way, the system ensures that the facemesh can capture all face points during the experiment.

Once these steps were completed, the explanation of both conditions of the experimental task (pro- and anti-saccades) was shown and the subject was informed whether she is facing an anti-saccades or a pro-saccade trial. The trial started with an empty circle representing a pro-saccade trial or a cross representing an anti-saccade trial, which appeared for a random time between 900 and 1500 ms. This cue was then replaced by a non-informative cue lasting 200 ms (an empty square in the center, same for both conditions). Then, a lateral stimulus represented by a full circle appeared for 150 ms. To avoid prioritizing visuospatial memory, boxes were drawn over the locations where the lateral stimuli would appear. The time allowed for the participant to respond was 650 ms and the ITI was 925 ms. Subjects had to complete a series of blocks on each of the tasks, alternating between the two task types (Figure 1D). Participants recalibrated the prototype between each block and the head position was checked (if not satisfied, the subject was asked to recalibrate). A complete run of the experiment included a tutorial block at the beginning with 10 trials of each type, then 160 anti-saccades and 160 pro-saccades, distributed in blocks of 20 repetitions each.

### Participants

26 participants completed a single session each (31.8 ± 6.1 years old, 7 women). All had normal or corrected to normal vision. The task was reviewed and approved by the IRB of Instituto de Investigaciones Médicas Alfredo Lanari (IDIM). All participants provided written informed consent in agreement with the Helsinki Declaration.

### Materials

We used the same computer configuration (Ubuntu 20.04.5 LTS / 16 GB RAM / Intel® Core™ i5-4460 CPU @ 3.20GHz × 4) and monitor (LG 22MP48HQ-P, 1920X1080, 60 Hz). We ran the experiments online and using the Logitech C270 (30 FPS, 720p) webcam.

### Analysis

#### Preprocessing

Raw data preprocessing was performed using pyxations (v 0.3.0), an open-source Python library for eye-tracking data in BIDS format. Specifically, pyxations handled (1) parsing of WebGazer CSV files into a per-sample DataFrame with trial-level metadata; (2) conversion of raw data to BIDS-ET directory structure via dataset_to_bids; and (3) propagation of trial-level behavioral metadata (e.g., saccade type, cue position, validation point coordinates) into the gaze samples files via the behavioral_columns parameter, eliminating the need for post-hoc CSV joins during analysis. Task-specific analysis (antisaccade error classification based on normalized gaze displacement and precision error metrics) was implemented on top of the pyxations output using standard Python data science libraries (pandas, numPy, scipy, statsmodels, matplotlib and seaborn).

First, the variability of the sampling frequency was addressed. This was done by resampling the estimates of each trial at 30 Hz using linear interpolation. We then performed a baseline subtraction of the average of the estimated x-coordinate during the fixation phase, defined as the median between -200 and 100 ms, to center it around 0, and then normalized it by setting the median between 500 and 700 ms as 1. The last 100 ms were excluded because sometimes participants made another saccade in anticipation of the next trial. Finally, trials with position values outside the range [-1.5, 1.5] were discarded: 6.9% ± 4.7% of pro-saccades and 8.4% ± 6.8% of anti-saccades. There were no significant differences in the number of accepted trials between conditions (Wilcoxon rank-sum test, p-value = 0.797). Thus, our preprocessing procedure did not introduce any bias in favor of one of the conditions.

Then we made a filter based on the number of errors in a block, if a participant made more than 50% of errors, we discarded the entire block. These extreme cases possibly correspond to a misunderstanding of the tasks in a given block. After using this filter, we discarded only 3 blocks in total (2 pro-saccade and 1 anti-saccade trial).

For better visualization of the curves, half of the trials were mirrored as if the lateral stimulus always appeared on the same side. The mirroring was done by multiplying the estimates of the trials in which the lateral stimulus appeared on the left by -1. Thus, all correct anti-saccades should show a jump in their estimates from a value close to zero to a negative value, whereas correct pro-saccades should show a jump toward positive values.

To quantify eye movements, we defined thresholds after normalization in the interval [-0.5 0.5], so that if the signal exceeded them (starting from the last range of our baseline, 100 ms), we considered that the subject was looking at one of the lateral targets. In this way, we can easily calculate the four possible categories (correct pro-saccades, incorrect pro-saccades, correct anti-saccades, and incorrect anti-saccades) and their corresponding times (RTs).

#### Statistical analysis

The first aim was to compare pro-saccades and anti-saccades performance. Thus, we performed planned comparisons between anti-saccades and pro-saccades: the percentage of correct responses and the RTs for correct responses. In both cases we used the nonparametric Wilcoxon rank-sum test. In a second level of analysis, we explored the anticipation effect in the anti-saccades by comparing the RTs of correct vs incorrect anti-saccade trials. In all cases, we report the median and interquartile range between participants, and the d.f. and p-value provided by the Wilcoxon test.

## Results

### Experiment 1: Accuracy assessment

We started evaluating our prototype in the accuracy assessment by measuring the sampling rate, and the global horizontal, vertical, and total errors (Table 1). The sampling rate was not constant along the task but very consistent across sessions, both in terms of mean values and variability (Table 1; see also top panels of figures 2A-B). Both participants presented a larger contribution to the total error in the vertical direction than in the horizontal direction (Table 1).

**Figure 2.**
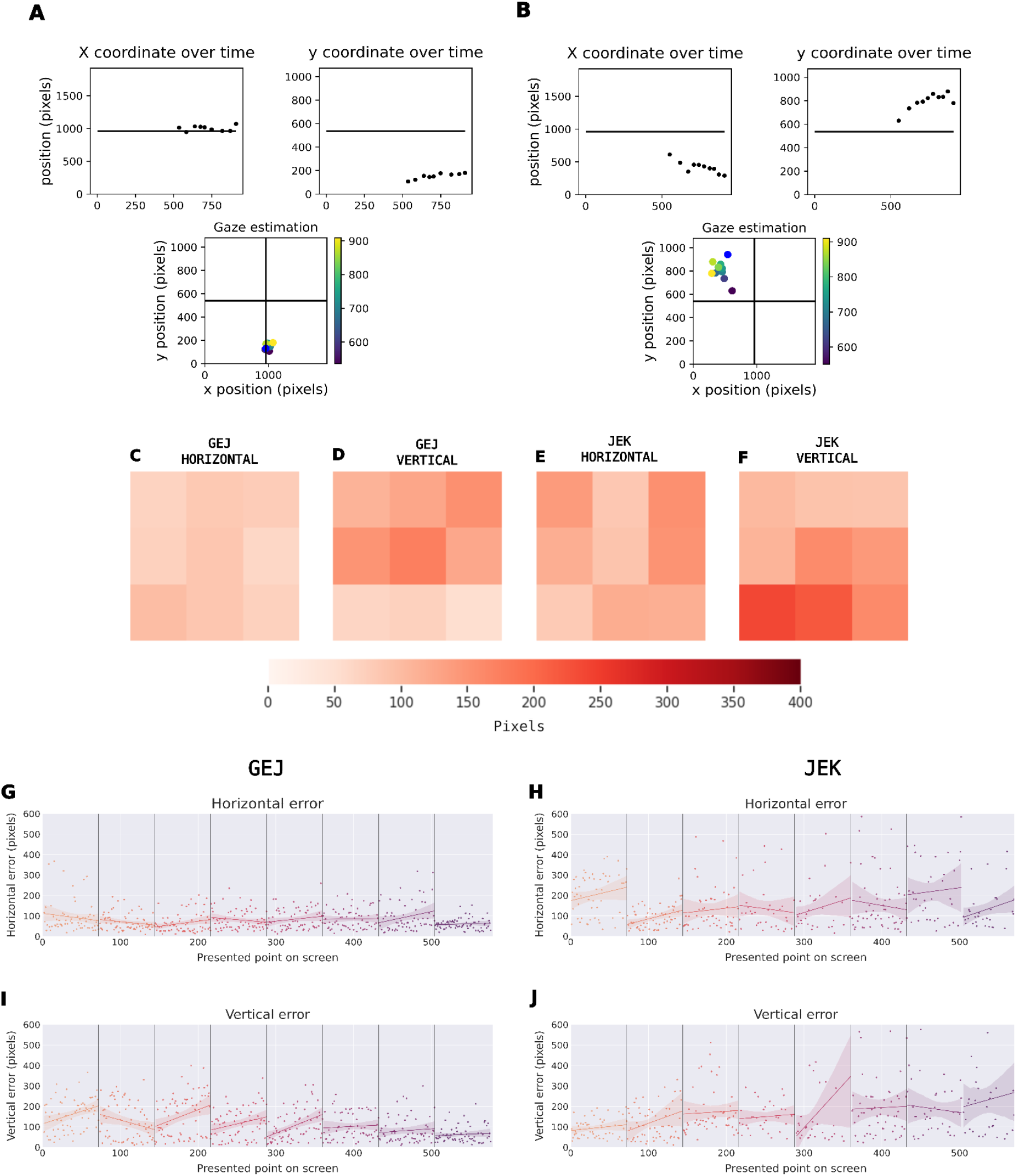
Accuracy assessment. ***A-B)*** Accuracy for one trial. The blue dot represents the circle presented on the screen. The upper graphs indicate the vertical and horizontal error time series and finally, the lower graph indicates the 2D spatial accuracy using the color scale for the time at which each eye tracker sample was acquired. Panel **A** shows a trial with low error and **B** shows one with higher error. ***C)*** Spatial error for a complete experimental session. The first graph corresponds to GEJ (left) and JEK (right). The colors of the map represent the error (vertical or horizontal, as indicated). ***D)*** Temporal error for each circle presented. The vertical lines represent the new calibration. The regression line is plotted for each round for display purposes only.

In general, the error values were smaller at the center and larger in the corners, both for horizontal (Fig. 2C,E) and vertical (Fig. 2D,F) errors. Finally, we decided to evaluate the reliability of errors in time by visualizing the average error at each validation round across the whole experiment. Figures 2G-J show that, except for a few noisier blocks, the error was roughly constant after each calibration (vertical lines) and across the whole experiment.

### Experiment 2: Anti-saccade task

Based on the capacities and limitations of the prototype and the relevance of the task, we used the anti-saccade task as a case study. In the task, participants have to fixate in the center and, based on a cue, they have to move their gaze toward or against a target that appears on the left or right side of the screen at middle height. Thus, the ET system has to determine whether they move their eyes to the correct side and the time they took to initiate the response. Therefore, the limitations on accuracy in space are not relevant because the regions of interest are only the center fixation and the hemispace, and a temporal resolution of 33-50 ms is enough to observe the difference in reaction times in this task.

After rejecting extreme trials and mirroring the left trial, the raw data can be observed in Fig 3A I, (only anti-saccades are shown), where the time 0 ms indicates the appearance of the target and 1 corresponds to the targets’ position. This data can be converted in degrees of visual angle using the output of the virtual chin-rest (Fig 3A II, see Methods section). As shown in Fig. 3A III, the vast majority of anti-saccade movements are against the target, i.e. position equal to -1 (correct responses) and some of them presented movements toward the target (incorrect responses). Although fewer, it is possible to observe that incorrect responses are faster than the correct responses when comparing the distribution of the response times (RTs) (top and bottom panels, Fig. 3A III).

**Figure 3.**
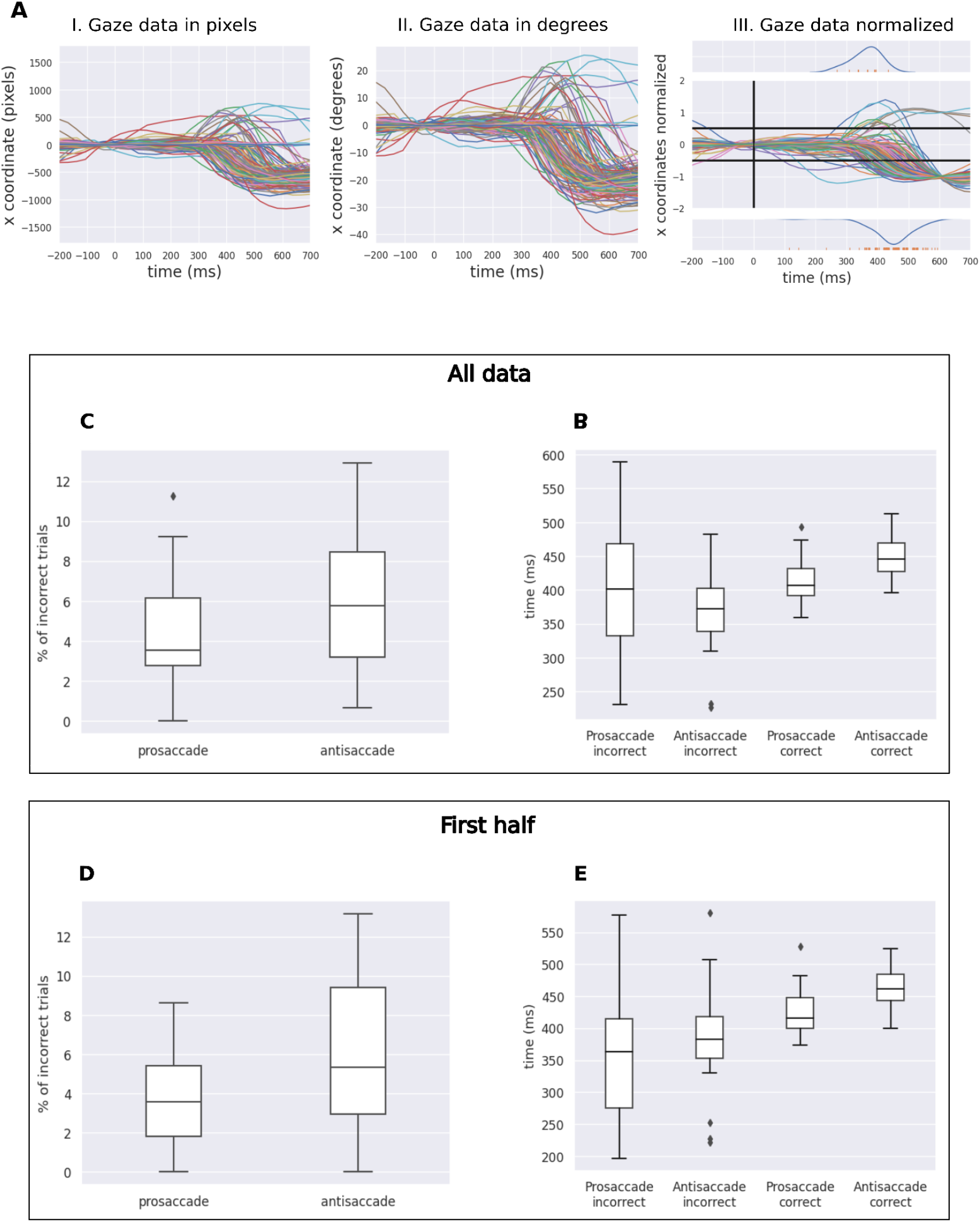
Anti-saccades performance. ***A)*** Pipeline Analysis for x-coordinate gaze prediction (one subject anti-saccade trials showed). Anti-saccades were mirrored within each trial to combine left and right saccades together. For details See *Preprocessing*. **I)** Data in pixels. **II)** Transformed data from pixels to visual degree angles using virtual chin-rest information. **III)** Minimum-maximum normalized data. Adjacent histograms on top and bottom of the gazes’ x-coordinate show the RT of errors and correct trials, respectively. The time 0 ms corresponds to the presentation of the lateralized cue, and the horizontal lines are the thresholds for saccades direction and the response time (RT). **B)** Error rates for pro-saccades and anti-saccades. **C)** Response times (RT) for correct and incorrect trials in pro-saccades and anti-saccades conditions. **D)** Error rates for pro-saccades and anti-saccades using only the first half of the experiment and excluding the first block. **E)** Response times (RT) for correct and incorrect trials in pro-saccades and anti-saccades conditions using only the first half of the experiment and excluding the first block.

As planned, we quantified the difference in error rates (Fig 3B) and RTs (Fig. 3C) for all pro-saccade and anti-saccade trials. Regarding error rates, the percentage of incorrect trials was higher for anti-saccade than pro-saccade trials, as reported previously in the literature (Currie et al. 1991; Hallett 1978; Hutton and Ettinger 2006; Munoz and Everling 2004). This difference did not reach significance at 0.05, but showed a clear tendency (median [IQR]; pro-saccades: 3.5 [2.8, 6.1] %; anti-saccades: 5.8 [3.2, 8.5] %; Wilcoxon rank-sum test (N=26): p = 0.087). In the same line, we found that in the correct anti-saccade trials the RTs were longer compared to pro-saccade trials (median [IQR]; correct pro-saccades: 416 [400, 448] ms; correct anti-saccades: 461 [444, 484] ms; Wilcoxon rank-sum test (N=26): p = 0.00019), as expected from the literature (Currie et al. 1991; Hallett 1978; Hutton and Ettinger 2006; Munoz and Everling 2004). This difference was significant and reflects the additional cognitive cost involved in the inhibition required to perform anti-saccade trials. Interestingly, we observed lower RTs for anti-saccades in incorrect trials compared to the correct ones (median [IQR]; incorrect anti-saccades RT 382 [353, 418] ms; correct anti-saccades RT 461 [444, 484] ms; Wilcoxon rank-sum test (N=26): p =2.0e-06), and also with the correct pro-saccade trials (median [IQR]; incorrect anti-saccades RT 382 [353, 418] ms; correct pro-saccades RT 416 [400, 448] ms; Wilcoxon rank-sum test (N=26): p = 0.0030). This was not the case in incorrect pro-saccade trials (median [IQR]; incorrect anti-saccades RT 382 [353, 418] ms; incorrect pro-saccades RT 363 [275, 414] ms; Wilcoxon rank-sum test (N=26): p = 0.24). This finding could reflect the presence of “fast errors” in this trial type, consistent with the idea that errors arise from a failure to inhibit automatic responses.

To understand the amount of data required to perform this task with our prototype, we analyzed the data again, using only the first half of the data (blocks 2-8), which corresponds to keeping 43.75% of the data, and around 10 minutes of task time without considering virtual chin-rest setup and calibration. As observed in Figures 3D and 3E, the results for half of the task were almost identical for incorrect trials (median [IQR]; correct pro-saccades: 3.6 [1.8, 5.4] %; correct anti-saccades: 5.3 [2.9, 9.4] %; Wilcoxon rank-sum test (N=26): p = 0.061) and RTs (median [IQR]; correct pro-saccades RT: 416 [400, 448] ms; and correct anti-saccades: 461 [444, 484] ms; Wilcoxon rank-sum test (N=26): p = 0.00019). Thus, acquiring around 10 minutes of data should be enough to reproduce lab results for this particular task.

Overall, after collecting the data from the anti-saccade experiment, we replicated a general result in the literature on this classic cognitive task: a higher number of errors were observed in anti-saccade trials (opposite to the stimulus) compared to pro-saccade trials (same direction as the stimulus).

## Discussion

Eye tracking (ET) is an established technology used in research, clinical assessment, and rehabilitation. In recent years, there have been some developments in using standard home computers and webcams to build eye-trackers for different purposes. These could be human-computer interactions, evaluation of digital platforms, or to conduct ET experiments over the web in an asynchronous and unassisted manner. Interestingly, several studies have shown that web-based ET settings have comparable data quality with respect to laboratory settings (Bridges et al. 2020; Germine et al. 2012; de Leeuw and Motz 2016; Semmelmann and Weigelt 2017). In this context, new algorithms are needed to deal with more noisy environments and more limited and non-dedicated hardware. In this study, we presented a web-based ET prototype and tested its reliability in two experiments: an accuracy assessment measuring spatial and temporal errors, and a classical cognitive experiment with clinical relevance, the anti-saccade task.

### Accuracy assessment

The web-based ET prototype presented here has an updated face-mesh model, an explicit calibration similar to in-lab setups, a head motion detection module, it incorporates a virtual chin-rest, and offered compatibility with the jsPsych library, as well as a browser-accessible interface for wider use. As a first step, we implemented an experiment to evaluate its performance. The accuracy assessment showed that our prototype had a relatively constant sampling rate. This is consistent with previous reports, although the specifications of the webcam, the processing power of the computer, the current computational load (such as the number of open browser tabs), and the inherent characteristics of the eye-tracking algorithm itself all determine the sampling rate upper limit (Bánki et al. 2022; Gagné and Franzen 2023; Semmelmann and Weigelt 2018). The spatial errors were higher in the vertical axis, as previously reported in other eye tracking experiments in online settings (Semmelmann and Weigelt 2018) and also in laboratory settings (Ke et al. 2013). It is more common for the gaze estimation error to be reported as a single value rather than disaggregated by direction. In this regard, total errors are comparable to those in previous literature (Heck, Becker, and Deutscher 2023; Papoutsaki et al. 2016). Moreover, the asymmetry of the errors should be taken into account when selecting experiments. For instance, our prototype is better suited for experiments where the horizontal axis is more relevant, such as the anti-saccade task. Also, the pattern of errors showed smaller magnitudes in the center and progressively larger values toward the corners of the screen. With the exception of a few noisy blocks, the error remained relatively constant after each calibration and persisted consistently throughout the experiment. Finally, by using the virtual chin-rest, we could measure indirectly the distance to the screen and obtain a robust estimate of the pixel-to-degree ratio, which was not assessed in most web-based ET studies.

### Validation of the remote eye-tracker using the anti-saccade task

Our results showed higher error rate and RTs in anti-saccade than pro-saccade trials. This is consistent with previous studies (Currie et al. 1991; Hallett 1978; Hutton and Ettinger 2006; Munoz and Everling 2004). Moreover, we showed that the errors corresponded to anticipatory saccades as they were faster than the correct responses. Our results fall within the expected range of variability for the percentage of antisaccade errors in healthy subjects reported in the literature. Specifically, our study found a median percentage of 5.8%, which is consistent with previous findings ranging from 2% to 25% (Reuter and Kathmann 2004). The fact that we found significantly longer reaction times in the anti-saccade than in the pro-saccade trials is consistent with the additional cost of inhibition present in the former condition. When analyzing incorrect trials, it is difficult to determine whether errors are due to an attentional component or some other underlying cause. However, when focusing only on correct trials, it becomes plausible that the observed time difference between conditions is due to an inhibitory process. Reflexive pro-saccades are rapid eye movements that occur automatically when we encounter novel objects, driven by the brain’s natural attentional focus on salient stimuli, also known as exogenous attention. These reflexive eye movements are fast and do not require conscious thought. In contrast, anti-saccades involve the conscious inhibition of the reflexive response and the voluntary redirection of attention, and thereafter gaze, away from the stimulus. Performing an anti-saccade requires inhibitory control to suppress the automatic pro-saccade and cognitive processes that generate the appropriate spatial coordinates for the anti-saccade (Olk and Kingstone 2003).

Interestingly, we found “fast errors” in the anti-saccade trials but not in the pro-saccade trials. This is consistent with the idea that the errors come from a failure on the inhibition of the automatic response (Ratcliff et al. 2016; Ratcliff and Rouder 1998; Roberts, Hager, and Heron 1994). To our knowledge, this effect was not described before for the anti-saccade task, and it would be very important to explore it in depth in future work as it could be a landmark of the processes involved. Studying more carefully eye movements in commonly used clinical cognitive tasks could provide us more specificity on the assessments (Linari et al. 2022).

The development of ET devices at a lower cost would make this field accessible to more researchers. In addition, the possibility of running ET devices that do not even require dedicated hardware opens up for potential applications in telemedicine. In this study, we explored this possibility using common webcams and presented a prototype that can be used in a real-life setting or remotely over the Internet. Some developments in this direction have been presented in recent years (Bánki et al. 2022; Papoutsaki et al. 2016; Semmelmann and Weigelt 2018; Xu et al. 2015). Although these methods have the advantage of a dramatically lower cost, they all have important limitations compared to a dedicated ET hardware. The most important is spatial resolution, which is about 4 degrees of visual angle compared to 0.5 in top-of-the-line ET equipment. Another important limitation is the sampling rate, which is given by the webcam, usually 30 Hz in most of the market models. Lower sampling rates limit the study of saccades (rapid ballistic eye movements) and fixations (the intervals where eye position is fixed, usually lower than 200 milliseconds). Notwithstanding, in these settings it is possible to estimate gaze position in tasks and paradigms where low spatial and relatively low temporal resolution are sufficient to classify responses and behavioral performance.

### Key contributions, limitations, and future work on web-based ET development

In this study, we made several key contributions to the development of a web-based ET prototype. First, we replaced the original face-mesh model used in WebGazer with an updated version. This improvement fixed a silent crash issue that made it unsuitable for experiments longer than 10 minutes and provided a more accurate estimate of eye position. Second, explicit calibration phases were implemented, replacing the reliance on continuous calibration through mouse responses. In this new type of calibration, users fixated their gaze on the calibration points while pressing the spacebar. This change improved calibration accuracy and reduced computational burden by eliminating the need for continuous calibration, while making it more similar to the calibration of traditional eye tracking devices. Third, a head motion detection module was incorporated to prompt recalibration based on eye positions and to mitigate the impact of head motion on gaze estimation. The prototype eye tracker also offered compatibility with the jsPsych library for experimental setups, as well as a browser-accessible interface for wider use. To facilitate exploration and development, minimal HTML and JavaScript playgrounds were created. In addition, we included the virtual chin-rest in the setup to control the distance to the screen and which adds the possibility to convert pixel distances to degrees of visual angle. Finally, the availability and usage details of the code were documented in a public repository on Github. Together, these contributions improved the accuracy, stability, and usability of the eye tracker prototype.

Even after these improvements, there are still several limitations. Firstly, the sampling rate is still variable and low for many purposes. In this work, we use linear interpolation to mitigate this problem, but this may be an issue for answering specific research questions that require a higher and consistent sampling rate. Secondly, although we tried to mitigate discrepancies of different versions of the hardware and software, for instance using the virtual chin-rest, this is an issue when developing web-based applications. Since Chrome is the most widely used browser in the world, we ran our experiments on this web browser. Nevertheless, it would be important to evaluate the experiments in some other popular browsers, such as Mozilla Firefox and Safari. Thirdly, the eye-blink detection is part of the pipeline of any top-of-the-line ET equipment, but it is not commonly included in web-based eye tracking setups for cognitive experiments. To our knowledge, only one paper estimated blinks based on web-based ET data (Madsen et al. 2021), by detecting peaks in the vertical gaze position after a 200-ms median filter, although these peaks may be generated by other artifacts, such as head movements, or others. Finally, another difficulty in moving ET experiments from the laboratory to the web is the low temporal and spatial resolutions (Heck et al. 2023) which we believe will be overcomed by the availability of better hardware (faster webcams or synchronizing smartphones) and browser capabilities (i.e., the webGPU API^9^ which exposes the capabilities of GPU hardware for the web (Kenwright 2022)).

We suggest that future studies should focus on some particular issues. First, this type of system doesn’t provide any information about the subject’s blinking behavior. The usefulness of blink detection is twofold: it allows a better calibration, which is used only in the presence of blinks, and it generates a richer data acquisition, which can be filtered based on this activity. Second, a deeper understanding of sampling rate variability and the factors that prevent the hardware from consistently performing at its maximum potential is needed. Third, different models should be explored to obtain a better estimation of the gaze. Webgazer uses a variant of ridge regression (Papoutsaki et al. 2016). However, given the modular nature of the software, it is possible to experiment with other models (e.g., neural networks). Additionally, based on the rapid advances in machine learning, it would be useful to try other models for obtaining facial landmarks and making predictions. The ongoing advancements in hardware optimization by web browsers, powered by technologies like WebGPU, hold great promise for enhancing the processing of webcam data and achieving faster inference times (Goh, Ho, and Abas 2023).

### Anti-saccades task: Diagnostic potential

Recently, multiple studies have highlighted the diagnostic potential of the anti-saccade task, particularly its ability to discriminate between Alzheimer’s disease (AD), mild cognitive impairment (MCI) and healthy populations (Boz et al. 2023; Opwonya, Doan, et al. 2022; Opwonya, Wang, et al. 2022). It is worth mentioning that, although we focused on the anti-saccade task, there are other tasks relevant to clinical research that do not require a high spatial resolution ET (Eizenman et al. 2003; Readman et al. 2021). Furthermore, ET can also be utilized to enhance the outcomes of gaze-contingent interventions, which provide online feedback based on eye movements in individuals with different brain disorders (Carelli et al. 2022). Based on this body of research, we believe that a fully functional web-based eye-tracking methodology could be extremely helpful in monitoring the progression of different pathologies, providing an opportunity for excluded populations with difficult or even no access to mental health care. Furthermore, it could serve as an early warning system, not only for diagnosis, but also for informing the health care professional of the patient’s condition, and only to make a medical visit if necessary.

## Conclusions

In conclusion, this study presented an open source, improved web-based eye-tracking prototype that is promising in terms of reliability and usability. It showed a relatively constant sampling rate and provided spatial errors comparable to previous studies. Moreover, the anti-saccade task, as a case study, showed higher error rates and longer reaction times for anti-saccades compared to pro-saccades, consistent with the existing literature. However, there are limitations to consider, such as low spatial and temporal resolutions, which could be partially addressed with advances in hardware and browser capabilities. Future work, such as blink detection and the evaluation of other gaze estimation models, is also an important part of this endeavor. Overall, web-based eye tracking holds promise for expanding research accessibility, reaching diverse populations, and potentially aiding in the diagnosis and monitoring of cognitive disorders. Further improvements and research are needed to increase the capabilities and accuracy of this technology.

## Declarations

### Ethics approval

The anti-saccade task was reviewed and approved by the IRB of Instituto de Investigaciones Médicas Alfredo Lanari (IDIM).

### Consent to participate

All participants provided written informed consent in agreement with the Helsinki Declaration.

### Availability of data and materials

Anonymised data for this manuscript can be found on GitHub: https://github.com/GEJ1/et_webcam_experiments

### Code availability

Details of implementation and usage are provided in the README file of the git repository of the prototype: https://github.com/ffigari/rastreador-ocular#usage. Analysis scripts for this manuscript can be found on GitHub: https://github.com/GEJ1/et_webcam_experiments.

### Authors’ Contributions

FF, GEJ and JEK designed the study; FF programmed the ET prototype; GEJ adapted the virtual-chin rest; GEJ collected the data; GEJ, AP and JEK analyzed the data; GJ, AP, and JK wrote the manuscript.

## AcknowledgmentS

We thank the contributions of Peter J. Kohler (Department of Psychology, York University) and Josh de Leeuw (Vassar College, New York) in the adaptation of the virtual chin-rest tool for jsPsych (https://www.jspsych.org/) (de Leeuw 2015).

## Footnotes

1 https://github.com/jspsych/jsPsych/discussions/2490

2 https://github.com/ffigari/WebGazer/compare/16f69474d40132c7faa826b2afc7fd464bc6c6c5..e5df9f9c3521ec3e384e962db49d94b2411789bb

3 https://github.com/brownhci/WebGazer/issues/171

4 https://github.com/ffigari/WebGazer/commit/16f69474d40132c7faa826b2afc7fd464bc6c6c5

5 https://github.com/ffigari/WebGazer/commit/7c6b7fb4dcefea2b85d7a24b3e86bd9a31b938d4

6 https://github.com/jspsych/jsPsych/pull/1442

7 https://www.jspsych.org/7.3/plugins/virtual-chinrest/

8 https://gs.statcounter.com/browser-market-share

9 https://developer.chrome.com/blog/webgpu-io2023/

